# From system-wide differential gene expression to perturbed regulatory factors: a combinatorial approach

**DOI:** 10.1101/013839

**Authors:** Gaurang Mahajan, Shekhar C. Mande

## Abstract

High-throughput experiments such as microarrays and deep sequencing provide large scale information on the pattern of gene expression, which undergoes extensive remodeling as the cell dynamically responds to varying environmental cues or has its function disrupted under pathological conditions. An important initial step in the systematic analysis and interpretation of genome-scale expression alteration involves identification of a set of perturbed transcriptional regulators whose differential activity can provide a proximate hypothesis to account for these transcriptomic changes. In the present work, we propose an unbiased and logically natural approach to transcription factor enrichment. It involves overlaying a list of experimentally determined differentially expressed genes on a background regulatory network coming from e.g. literature curation or computational motif scanning, and identifying that *subset* of regulators whose *aggregated target set* best discriminates between the altered and the unaffected genes. In other words, our methodology entails testing of all possible regulatory *subnetworks*, rather than just the target sets of *individual* regulators as is followed in most standard approaches. We have proposed an iterative search method to efficiently find such a combination, and benchmarked it on *E. coli* microarray and regulatory network data available in the public domain. Comparative analysis carried out on simulated differential expression profiles, as well as empirical factor overexpression data for *M. tuberculosis*, shows that our methodology provides marked improvement in accuracy of regulatory inference relative to the standard method that involves evaluating factor enrichment in an individual manner.

## Background

The availability of high-throughput technologies in recent years has made it possible to track the dynamics of a cell’s functional organization on the whole genome level [1,2]. The profile of genome-wide expression together with knowledge of the heterogeneous network of intricate molecular interactions determines the cellular phenotype, and holds the potential for providing insights into how a cell adaptively responds to environmental cues [2-5]. Functional genomics experiments such as those based on microarrays or RNA deep sequencing are a rich source of information about the cellular milieu, and provide a starting point for generating causative hypotheses about biological mechanisms [6-9]. A question that routinely needs to be addressed in large scale expression studies is the identification of key regulatory pathways underpinning coexpressed, or differentially expressed genes. Transcription rates are controlled in part by a complex network of regulatory interactions involving DNA-binding transcription factors (TFs) and *cis/trans* acting regulatory sequences distributed throughout the genome [10-13]. Changes in the functional activity or expression of one or more of these proximally acting regulatory proteins – possibly representing consequences of signaling events initiated farther upstream – can directly cause reshaping of the transcriptome. The inference of this set of ‘perturbed’ regulators is an initial and important step towards arriving at a broader mechanistic interpretation of altered genome-scale expression.

*De novo* approaches seeking to discover shared factor-binding DNA sequence motifs within the promoter regions of altered genes provide a rational starting point [14,15]. Such regulatory information for the genomes of many species continues to accumulate at a rapid rate from ChIP-seq experiments as well as low-throughput studies [16-23]. Databases providing experimentally determined/predicted transcription factor binding sites, TF motif profiles, and even meta-network information curated from literature evidence [24-31] are routinely available now, and could be usefully exploited by experimentalists interested in understanding differential TF activation in specific contexts. Towards this end, many bioinformatics tools have come up in recent times that facilitate such regulatory analysis. These methods [32-43] share the common denominator that an input list of genes specified by the user, e.g. coming from a microarray study, is overlaid on a prespecified background regulatory map, which may have been put together by combining information from diverse sources. In order to deal with the noisy nature of the data, some appropriate statistical test is applied to each TF in the back-end network to determine a statistically significant association, or over-abundance, between the targets of the TF and the input gene list, relative to the overall genomic background. Depending on the over-representation p-values computed, a prioritized list of candidate regulatory factors likely to be most relevant for interpretation of the user’s data is thereby generated.

A few examples of such applications are noted here. ChIP Enrichment Analysis (ChEA) is one such popular tool that leverages a curated database of ChIP-seq profiles from mouse and human experiments to compute over-represented target sets via Fisher’s exact test of significance [32,33]. Two related applications, Kinase Enrichment Analysis (KEA) and Expression2Kinases (X2K), are methodologically similar but go a step further and, by additionally exploiting curated data on kinase-substrate relationships, suggest signaling pathways highlighted by input lists of altered genes [34,35]. ENCODE ChIP-Seq Significance Tool is a web-based interface which allows users to mine a back-end comprised of mouse and human TF binding site data generated as part of the ENCODE series of experiments [36]. Hypergeometric test is applied to score individual transcriptional regulators for significant association with the input list of genes. This test is similarly the basis for TF enrichment analysis implemented within the RENATO [37] and WebGestalt [38] tools. Other utilities such as Whole-Genome rVISTA [39,40], Promoter Integration in Microarray Analysis (PRIMA) [41], Cis-eLement OVERrepresentation (Clover) [42] and Relative OVER-abundance of cis-elements (ROVER) [43] work instead with the binding site motifs of known TFs, represented as position weight matrices (PWMs), information about which can be found compiled in resources such as TRANSFAC, JASPAR, HOCOMOCO, UniPROBE etc. [27-31]. Despite differing in the actual criterion applied for assigning target genes to every regulator, which is based on scanning of promoter sequences for high-scoring motif matches, they all nonetheless follow the common theme that over-abundance scores relative to the genomic background (i.e. p-values) are calculated for each regulatory motif *separately* against the *entire* list of input genes. Moreover, the null background implicitly assumed in all the above approaches is essentially one of no association, corresponding to a random distribution of altered genes over the genome.

When evaluating individual TFs for association with a large gene set detected as differentially expressed in a transcriptomic experiment above some user-defined threshold for significance [44-46], it is worth noting that the above methods are likely to work well when one or only a very small number of TFs have been differentially activated. On the other hand, if a gene list represents the collective consequence of perturbing multiple regulators, then it is conceivable that individual TFs may well fail to show up as statistically significant on application of one of the tests previously mentioned. The hypothetical situation in Figure 1 serves to illustrate this point. It is not hard to imagine a case where the target set of factor A, or B or C by itself may not show statistical association with the full input gene set I. The effect of other regulators acting concomitantly raises the possibility that any one TF may fall short of achieving separation between the altered and unaltered genes, when assessed in terms of the corresponding p-value of enrichment. Thus, the deductions about relevance of individual transcription factors for alterations in gene expression might be inaccurate.

From Figure 1 it also follows that aggregating the targets of A, B and C together would yield a more significant association of this *union* with the differentially expressed gene set. This suggests an alternative approach to delineating a set of immediately upstream transcriptional regulators causally underlying an altered expression profile. In the present work we propose a perspective which entails the testing of all possible *subsets* of TFs, i.e. the unions of their target genes, instead of just the target sets of individual TFs. In our opinion, this reformulation would appear to be a more natural and powerful approach to ascribing differential gene regulation to a set of perturbed transcription factors. A hypothesis for differential expression is therefore proposed by identifying the TF *combination* that collectively best separates the differentially regulated genes from the unaltered genes derived from any high-throughput expression profiling experiment.

## Results

### The default approach can lead to inaccurate inferences in an 'idealized’setting

A microarray expression profile can be regarded as representing a combination of signal (S), the ‘true’ pattern of differential transcription resulting from altered TF activity, and noise (N) introduced by various sources which result in the occurrence of false positives/negatives. We begin our analysis by considering TF enrichment in the N → 0 deterministic limit, where all the gene targets of a randomly chosen group of TFs are labeled as differentially expressed. This noiseless limit can be expected to provide an upper bound on the efficacy of any statistical enrichment test for regulatory inferences. We first assess the *default*, or standard, method of testing which involves estimating over-abundance p-values for *individual* TFs. What needs to be evaluated is how the p-value of every TF varies with the number of regulators present in the input set, and the possible effects of target set size and TF-TF overlap/co-targeting of common genes on the ascribing of statistical significance.

Figures 2 and 3 summarize the results for 10^4^ runs with simulated expression data on the *E. coli* RegulonDB network [26]. The efficacy of the default method has been gauged in terms of the frequency distribution of TF occurrences in the enriched set. This is compared with the underlying 'input' TF distribution, which to the first approximation is uniform. Figure 2 displays the comparison for a p-value significance threshold of 0.05, with TFs being ordered according to their out-degrees along the horizontal axis. Two features that Figure 2(B) illustrates are the overrepresentation of the higher degree regulators in the output set on the one hand, and a dip in the frequencies of occurrence of the TFs with small target sets on the other. This can be seen by a direct comparison of the input (blue) and enriched (red) distributions. Similar trend is also seen for other choices of the p-value threshold, in the range 0.01-1e-6. These features suggest that when multiple TFs are simultaneously perturbed, regulators with smaller target set sizes might fail to show statistically significant enrichment for the combined differentially transcribed gene set. Further, TFs which are themselves not differentially activated, but co-target genes with other regulators, can show a spurious association when evaluated by the hypergeometric/Fisher’s exact test.

Figure 3 presents the above trend from a slightly different angle. The percentage of runs in which any given regulator is included in the input set, but does *not* show a significant overabundance, is plotted against the ordered sequence of TFs ranked by increasing out-degree. This representation is consistent with the dip at the left end seen in Figure 2B, and brings out the possibility that, when evaluated one at a time against the combined set of *all* differentially expressed genes, TFs with smaller target sets might be missed out by the test for significance when regulatory activity of multiple factors is altered in concert. Thus, an artefactual under-representation

### Benchmarking reveals the best among three iterative search methods

We now explore testing combinations of TFs for association against the differentially expressed genes, instead of treating each TF separately, such that the combination which is found most consistent (in terms of minimum collective p-value for association) provides an alternative hypothesis for the altered regulatory activity. However, the infeasibility of exhaustively searching the space of possible combinations presents a practical stumbling block. For example, in typical bacterial networks, there are around O(100) transcription factors [26,69,70]. With the numbers of differentially expressed genes that usually occur in genome-wide datasets, the number of overlapping TFs would be of the same order of magnitude, requiring the computation of O(2^100^) or ~ O(10^30^) enrichment p-values. Even if we restrict the biologically sensible combinations to involve not more than 20-30 TFs, this would still leave about 100!/(30!*70!) possibilities to be tested, and even this number would be beyond the limitations of computational tractability. With eukaryotic gene regulatory networks and especially those for higher organisms like mouse or human, a factor of 10 increase over prokaryotes in the number of regulators to O(500-1000) is expected, based on standard regulatory meta-network datasets used in the literature [19-25]. This would translate into an even larger number of candidate combinations that would need to be swept over in order to identify the best-fit hypothesis. We therefore seek a solution that is sufficiently close to the global minimum, through a computationally efficient heuristic approach.

As the nonlinearity of the objective function sought to be minimized precludes the adoption of exact linear programming methods, we have tested three iterative search methods to efficiently arrive at such an approximate solution, which are described in the *Materials and Methods* section. Method A (Figure 4) is linear in the dimensionality of the search space, i.e. in the number of TFs tested, and the solution is built up by sequentially adding TFs in increasing order of their individual p-values. In the case of the Method B (Figure 5), it is easy to see that O(N) p-values have to be estimated in every iteration, because the differentially expressed gene set shrinks at every step as the genes already covered by the TFs selected up to that point are systematically eliminated from consideration. The origin of a quadratic execution timescale follows from this. The O(N^2^) scaling of Method C (Figure 6) also derives from the fact that the p-values corresponding to the TFs not already included in the growing solution have to be recomputed at every update.

We note that, even though the two O(N^2^) approximations proposed here can be expected to yield better results than the preceding linear-time greedy search, they still represent only local sampling of the search space, and so, like Method A, cannot ensure that the minimum found represents the true exact solution.

Figure 7 displays the results for the combinations obtained by running each of the three heuristics, Methods A-C, on the 7 GEO datasets [52-55]. This selection is not intended to be exhaustive in any way, but is representative of the microarray data available in the public domain, and serves to illustrate the salient features of our approach as well as some statistical properties of the search space. It is seen clearly that across all the 7 conditions analyzed, Method C outperforms the other two heuristics, yielding solutions with lower enrichment p-values. It is also observed that, consistent with our expectations, the O(N) Method A is found to be least effective of the three methods.

A three-way comparison was also carried out on the larger M3D *E. coli* expression dataset [57]. Based on the set of perturbed genes identified in each condition, the associated upstream transcription factors were then inferred by applying the three approximations in turn. The results of this analysis are summarized in Table 1. We assessed the performance of Method C separately against Methods A and B. In line with the outcomes in Figure 7, we find that Method C displays improved performance overall. For instance, in over 98% of the experiments, Method C yields a significance p-value that is at least as low as the outcome of Method B. Similarly, in no experiment does Method A improve on the p-value yielded by the application of Method C. It may also be noted that these results are reproduced across three different choices for the Z-score threshold (2, 2.5 and 3).

**Table 1.**
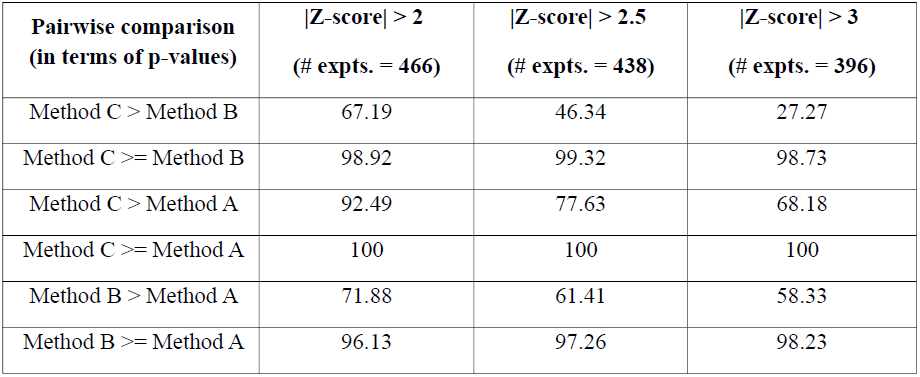
Benchmarking iterative search methods on M3D expression profiles. Three-way comparison among the heuristics proposed, based on 466 experiments with *E. coli* compiled under M3D. Each column displays the results for a particular choice of the Z-score cutoff applied for identifying genes with altered expression in every experiment. Numbers in brackets in the column headers are counts of those experiments in which at least one altered gene is targeted by a TF from the underlying network. All comparisons are percentages relative to the corresponding total number of experiments being considered. Performance of each method on every expression profile is quantified in terms of the over-representation p-value for the TF combination it converges to.

Out of the 466 experiments in total, with a Z-score cutoff of 2.0, we found 68 conditions in which the differentially transcribed gene set sizes were small enough such that only ≤ 14 TFs were found to overlap with these genes. In these instances, it was possible to identify the exact solution by enumeration over all possible combinations, allowing for a direct comparison with the results of the greedy methods. Across these 68 cases, we have estimated the number of conditions in which each of the three methods yields a p-value equal to that of the globally top-scoring combination. In addition, the quality of the obtained approximate solution has been assessed by its rank in an ordered list of TF subsets representing the full search space. Figure 8 summarizes the results of this exercise, supporting our earlier conclusion about the better performance of the O(N^2^) Method C, which not only yields the true best p-value more often but is also able to get closer to the global minimum, as assessed by the average rank of the obtained solutions. Similar results were obtained with the more stringent Z-score threshold of 3.0 as well, displayed in Figure 9.

### Search space can be rugged with multiple local solutions

The greedy heuristics considered above represent incremental approaches in which only the local neighborhood – comprising the current solution and its neighboring combinations differing at most by a single TF – gets sampled at every update. When multiple local minima coexist [58,62], local search might only yield a sub-optimal solution. In such a complex search space, stochastic approaches like simulated annealing (SA) or genetic algorithms with a larger radius of convergence might be preferable [61,62]. However, even these latter approaches do not guarantee convergence to the global extremum in a single run, and it is a known fact that SA requires fine tuning of the operational parameters for optimal performance, which is problem-dependent [63]. It is therefore of relevance to understand, to what extent an iterative search can achieve a favourable balance between speed (local methods are typically faster) and quality of the obtained solution (which is generally better in a suitable optimized SA procedure).

Greedy searches like the ones investigated here may not provide any benefit over simple steepest gradient method in a solution space which is smooth containing one or only a few local minima. In order to address this possibility, we have attempted to estimate how rugged the solution space is [58-60]. Towards this end, we implemented gradient descent search starting from different randomly drawn initial configurations (TF subsets), tracking the number of local minima attained as a function of the number of runs (starting configurations sampled). Figures 10-16 depict the results of this analysis for the 7 microarray datasets on which we ran the iterative procedure earlier. Over 2000 runs, the search is found to converge to multiple local minima, suggesting that the landscapes, at least for these specific examples, are moderately rugged. In such a rugged search space, not only does steepest gradient search have to be run repeatedly with random restarts to sample the multiple local minima, but even a large number of runs cannot guarantee that all the minima, and in particular the global minimum, would be covered. The gradient descent method as implemented here is therefore slow and in general inefficient.

The above summary statistics suggesting rugged solution spaces was further used for assessing the performance of the approximate Method C discussed earlier. Although the time complexity of both Method C and gradient descent is the same and O(N^2^), Method C is deterministic in the sense that the initial combination is not selected arbitrarily, so a direct comparison between the two for a single run of gradient descent search would depend on the choice of initial condition for the latter. A more sensible comparison would be to ask, in what proportion of runs of steepest descent (with random sampling of the starting combination) the converged solution represents an improvement over the result of Method C. The results of this comparison are tabulated in columns 3 and 4 in Table 2. We have compared the p-value following from application of Method C with the sorted list of p-values (in increasing order) yielded by gradient search with random restarts. Column 4 indicates that in 4 out of the 7 conditions, Method C attains the top-ranked p-value identified by gradient search method. Even in the case of the heat shock dataset where the rank is somewhat lower (62/814), the solution yielded by Method C still appears within the top 10% of the ordered list.

Considering the moderate ruggedness of the search space besides the broad range of differences in the objective function values among the various local minima, we also implemented a simple version of simulated annealing [61-65], which involved carrying out multiple independent runs of an exponential annealing schedule with different settings for the rate parameter *r*. Figure 17 illustrates the evolution of the p-value over one run of SA applied to the pH 5.0 dataset, and the solution obtained by Method C is additionally shown for comparison. In columns 5-7 of Table 2, the minimum p-values yielded by the SA procedure have been tabulated against the corresponding results from Method C and gradient descent method. SA provides an independent objective benchmark against which to gauge the outputs of the heuristic methods. Making the reasonable assumption that the solution following from SA represents the true global minimum, our analysis summarized by columns 5-7 shows that the approximate method C converges to the top-ranked solution in 4 of the 7 examples, and gets close - up to second best – in all but one of them (the heat shock profile). A similar comparison was also made on the M3D dataset [57]. The Z-score profiles were sorted according to the number of TFs targeting the significantly altered gene set, and the two methods were then applied to the subset of top 20 profiles, for which the search space dimensionality (number of overlapping TFs) ranged between 110 and 140 TFs. In 11 (55%) cases, the iterative Method C yields p-values which matched those obtained by the SA implementation. We would like to point out that these comparisons need to be assessed additionally keeping in mind the relative differences in execution speed as well, because for practical implementation achieving a trade-off becomes important. Both the gradient search with restarts and our implementation of SA require multiple runs with different starting states and/or settings, and *each* run involves O(N^2^) p-value computations, equivalent to the execution time of Method C. The number of runs of gradient search required to adequately sample the search space is expected to grow with its dimensionality (N), and thus become even more of a practical drawback in the larger eukaryotic TF networks. The examples we have considered here, although not exhaustive, make a case for Method C which is able to arrive at a good solution efficiently. Taken together, the results summarized by Figures 9 and 10 and Table 2 provide support for the efficacy of Method C in determining the maximally discriminative combination of TFs from a binarized profile of gene expression changes. This method should be particularly useful when data from elaborate expression studies involving large numbers of conditions/time points need to be analyzed, where stochastic search would be considerably slower.

**Table 2.** Summary of results for seven GEO microarray datasets. Benchmarking of Method C v/s gradient descent search based on 7 microarray datasets describing *E. coli* stress adaptation. The latter has been run 2000 times with random restarts for each dataset. Also tabulated are the p-values obtained from the simulated annealing (SA) procedure, which provides an independent estimate of the global minimum.

| Experiment | Number (resp. %) of genes found differentially expressed ($p < 0.001$, log fold change $> 1$) | % of runs of steepest descent search yielding a local solution at least as good as the output of Method C | Rank of solution found by Method C in a sorted list of local minima sampled by steepest descent search | Method C output log p-value | Minimum log p-value from steepest descent search | Minimum log p-value from SA |
| --- | --- | --- | --- | --- | --- | --- |
| pH 5.0 | 176/4345 (4.05%) | 6.1 | 1/251 | -43.4447 | -43.4447 | -43.4447 |
| pH 8.7 | 92/4345 (2.12%) | 57.6 | 1/16 | -35.7941 | -35.7941 | -35.7941 |
| Norflox | 312/4345 (7.18%) | 2.9 | 1/55 | -19.373 | -19.373 | -19.373 |
| Sucrose stress | 277/4070 (6.81%) | 0.8 | 1/37 | -22.1718 | -22.1718 | -22.1718 |
| NaCl stress | 194/4070 (4.76%) | 27.25 | 2/142 | -24.2429 | -24.3237 | -24.3237 |
| Stationary phase | 2324/4493 (51.72%) | 0.05 | 2/356 | -32.0485 | -32.5571 | -32.5571 |
| Heat shock | 3146/4493 (70.02%) | 5.26 | 62/814 | -43.8734 | -47.2333 | -47.3869 |

### The proposed approach leads to less biased, more accurate inferences of differentially acting TFs

We now return to the idealized setting described earlier in *Results*, with random combinations of TFs being selected for perturbation and used as a basis for assigning genes to altered/unaffected sets in a deterministic manner. We found that when statistical enrichment is assessed for single TFs against the full set of differentially expressed genes, the sub-dominant TFs with smaller target sets get under-represented in the recovered set (based on adjusted p-values ≤ some threshold), while the global regulators tend to occur more often than they are actually selected for inclusion in the input set (Figures 2 and 3). We would like to compare this outcome with the results of the Method C which provides an approximation for the TF combination with least collective p-value. Figure 18 displays the result for the distribution of TF occurrence frequencies across 10^4^ random trials. This plot is the same as in Figure 2, except that now the results for Method C have been additionally superimposed (in blue). Our alternative approach reduces the underrepresentation at the low degree end, and at the same time also alleviates the over-occurrence of the high degree TFs. This improvement is quantified in terms of the root mean-squared error (RMSE), which shows a > 2.5-fold decrease (0.0368 for Method C against 0.0954 for the default method).

Overall, the degree-dependent bias suggested by the red curve (the standard approach of testing single TFs) is to a fair extent suppressed in our method.

We find that the strategy of identifying the top-ranked combination of TFs also leads to improved accuracy in the recovery of TFs, i.e. it recapitulates the input set of TFs better. This is seen from Figure 19, which displays a two-dimensional scatter plot of the accuracy value pairs (default v/s Method C) for every single trial, at p < 0.05 threshold for the default method. Points lying above the 45 degree diagonal represent trials in which Method C recovers a better solution set than the default approach, and vice versa. For the RegulonDB graph [26], in 94.7% of the trials Method C achieves improved performance compared with the default methodology (as opposed to only 2.6% going in the opposite direction), and yields significantly higher accuracy values overall (Wilcoxon signed-rank test [66] p-value = 0). This result is fairly insensitive to the choice of the cutoff for assigning statistical significance in the standard test, and is reproduced over a range of threshold values (0.01-1e-6).

Revising the above sampling procedure by now selecting TFs in accordance with the TF-TF interaction structure of the underlying graph, we find that this modification does not change the basic result obtained previously. As displayed in Figure 20, Method C still shows improved accuracy for recovering the input subset of TFs in comparison to the standard methodology of evaluating each TF separately. At p = 0.05 cutoff, Method C yields higher accuracies compared to the default approach in 85.4% of the trials, as opposed to only 7.43% of trials showing the reverse trend.

A second modification we considered is the addition of noise, mimicked by introduction of a small proportion of misclassified genes, wherein 5% of genes are randomly selected from the network and reassigned to the opposite category (differentially expressed v/s unaltered). As shown by the profile of occurrence frequencies in Figure 21, the deviations from the nearly flat input distribution are largely suppressed upon application of Method C. For example, at p < 0.05 cutoff, the RMSE decreases from 0.0796 (standard approach) to 0.0211 (Method C). Pair-wise comparison of the accuracy values of the recovered TF sets, displayed as a two-dimensional scatter plot in Figure 22, shows that in 85.8% of trials Method C improves on the output of the default significance test, and also yields two-fold increase in the number of trials in which unit accuracy is obtained (603 v/s 286, out of 10^4^ trials). Thus, once again we find that the earlier trends are essentially reproduced here, and it is evident that a small distortion of the idealized differential expression pattern has minimal impact on the validity of the earlier result. Taken together, the preceding results make a case for the utility of the combinational approach as an improved and logically sensible alternative for generating less biased, biologically meaningful hypotheses for altered TF activity from global expression data.

### The combinatorial approach applied to *M. tuberculosis* TFOE data demonstrates marked improvement in recovery of the causal TF

As another independent validation of the methodology presented here, we applied both the standard approach and the combinatorial search to microarray profiles for *M. tuberculosis* coming from recently published TFOE experiments [67,68], to see how well the network based inference alone (disregarding expression change) can recover the causal upstream TF. Out of 78 TFOE experiments where the over-expressed TF was present in the reference regulatory network, we noted that in only 48 cases the target gene set of the TF had a non-zero overlap with the differentially expressed gene set. This is merely indicative of the incomplete nature of the background network used [69,70], and illustrative of a limitation which can be expected more generally when carrying out system-level analysis, on which scale the availability of interaction information is unlikely to be complete and altogether accurate. Our comparison shows that the evaluation of each TF separately by Fisher’s exact test is able to identify the causal TF in only 21 (44%) cases (at the maximum acceptable p = 0.05 cutoff); in contrast, the TF combination arrived at by Method C contains the causal TF in 32 (67%) cases, representing ≈ 50% improvement. In fact, there are 16 experiments in which Method C uniquely identified the over-expressed TF, which by itself fails to show significant enrichment when tested against the full set of differentially expressed genes (this may be contrasted against only 5 cases where the opposite is true). We further wanted to assess whether a simple ranking of the individual over-abundance p-values, regardless of whether they are deemed significant or not, is able to reveal the correct TF. In none of the 16 cases mentioned above, the over-expressed TF is found ranked first in the sorted list. It is interesting to additionally observe that in 10 of the 16 cases, not only is the causal TF not over-represented individually, but it even fails to show up among the top *k* sorted TFs, where *k* denotes the size of the corresponding non-redundant TF combination identified by Method C. These differences in outcome once again underscore the limitation of assessing TFs one at a time, even in application to single TF OE profiles, with scope for improvement being suggested by the combinatorial perspective explored in the present study.

## Discussion

Rewiring of transcriptional regulatory networks under different perturbations is an important problem that needs to be understood on the systems level. Several methodologies have been proposed in the literature to interpret data derived from high-throughput profiling experiments in terms of enriched biological functions or over-abundant proximal regulatory elements [71-75]. The particular approach which has been the starting point for the present work involves reducing the case v/s control comparison of large scale gene expression to a binary profile with a step-like threshold, where each gene is classified as either differentially expressed or unchanged according to the result of some suitable chosen statistical test applied to the gene expression values. Pathways or DNA *cis*-regulatory motifs which over-occur in the differentially expressed subset are then inferred based on the null hypothesis that the significantly altered genes are randomly distributed over the genome and have no statistical association with the functional gene set under consideration. We note that the results of this approach are dependent on the threshold deemed significant for calling a gene differentially expressed, and would in general vary as the dichotomous assignment changes according to different choices of this confidence p-value. Other methods proposed in the literature work instead with the unfiltered expression values directly and are less sensitive to this subjectivity. For example, the popular methodology GSEA [75] computes enrichment scores for gene sets (such as pathways or targets of regulatory motifs) starting with an ordered list of all genes ranked according to some measure of expression change (e.g. t-test score), to identify gene sets whose members show concordant changes in expression between two phenotypes. Furthermore, GSEA and its related offshoots [75-77] use a permutation-based approach of randomly reassigning sample labels to generate a null distribution of scores, which thus differs from the negative control defined for the hypergeometric test that assumes a random binomial distribution of gene labels to gauge statistical association. While Fisher’s exact test applied to binarized data, which is the basis for the present approach, has its limitations, this sort of test is in fact used quite widely in enrichment analyses, as illustrated by the examples mentioned in the introductory section [27-43,74]. Thus, the motivation for the present work is quite justified, and the relevance and utility of the alternative approach we have proposed and explored here should be assessed against the backdrop of these other currently existing tools for transcription factor analysis, all of which are essentially based on application of either the hypergeometric test or some close variant thereof.

It is suggested here that an equally acceptable hypothesis for differential regulation can instead be arrived at by seeking a group of TFs which is *collectively* most predictive for the altered large-scale expression. This approach identifies a candidate set of TFs that is overall distinct and *not* trivially obtainable from a sorted list of TFs arranged in ascending order of their *individual* over-representation p-values. As this alternative methodology presents the practical difficulty of having to deal with a solution space whose size grows exponentially with the overall number of TFs in the network, greedy heuristics were proposed and their efficacies compared. Our comparison between two quadratic-time methods for estimating an approximate solution (based on application to *E. coli* microarray data), in particular, holds some relevance going beyond the problem studied here. Many machine learning approaches to classification, e.g. logistic regression, involve identifying a small subset of discriminative features from a high-dimensional, more diverse feature vector. Linear methods based on sequentially adding features to a growing set based on the discriminative performance of each individual feature are usually adopted for this purpose (e.g. [78]). Our analysis of Method C suggests an alternative take on feature selection which might lead to improved classifier performance, especially when dealing with a large number of potential features. We reiterate that this procedure is distinct from the other iterative method (Method B) which is analogous to the set cover heuristic [50].

A possible direction in which the current work could be extended would be to incorporate the principle of parsimony. From the point of view of practicality, a simpler hypothesis (i.e. a smaller number of inferred TFs) might be more favorable, and so, instead of the global *minimum*, an *optimal* subnetwork could be sought which strikes a judicious balance between the TF set size and the fit to the data (over-abundance p-value), analogous to Bayesian/Akaike information criterion for parameter estimation in multivariate regression [79]. This could be implemented in the iterative procedure by, e.g., requiring that every additional TF incorporated into the growing solution provide some minimum reduction in the overall p-value. Thus, once the point is reached beyond which further addition of TFs does not produce substantial improvement, the algorithm could be terminated. Since we have wanted to keep the analysis presented here fairly general, this additional constraint has not been imposed, as it would require introducing a user-defined parameter which decides the balance between parsimony and fit, and the outcome of the procedure would then depend on the numerical value of this additional parameter.

Finally, we note that the present methodology could be revised to incorporate combinatorial effects of TFs, especially relevant in the case of eukaryotes and higher organisms where cooperative synergistic interactions between pairs (or even larger groups) of TFs are quite common in the transcriptional control of gene expression [80-83]. In particular, if a TF requires co-binding of other regulators to nearby DNA sequences to modulate the rate of transcription of its gene targets, then an analysis based on the entire set of binding sites of that TF, as identified e.g. from enriched peaks in a ChIP-seq genome-wide binding profile [16], may not give correct results, as many of those binding sites may not translate into bona fide causal regulatory interactions. Thus, the use of gene sets which are co-targeted by multiple TFs (i.e. regulatory ‘modules’) as building blocks, instead of the target sets of individual regulatory motifs as has been followed here, might provide more realistic hypotheses for discriminative regulatory subnetworks. The results of such an analysis could in fact even be leveraged to reveal novel TF-TF combinatorial effects. These ideas are currently under investigation.

In summary, we have proposed a general, unbiased, and logically natural methodology to come up with a set of differentially active regulatory factors implicated by large-scale differential gene expression data, which avoids the somewhat artificial breaking up of the problem that happens when testing each regulator separately. By effectively boosting regulators with smaller target sets, our strategy holds out the possibility of revealing biologically important regulators which might be missed by many standard approaches that all share the common feature of assessing each regulator separately for overlap against the full list of altered genes. This perspective should be of immense utility in contributing to a clearer mechanistic understanding of the global transcriptional remodeling underpinning adaptation, or dysfunction.

## Methods

### Re-examining the default approach to identifying perturbed regulators in the noiseless limit with simulated profiles

We assume a deterministic setting in which a subset of transcription factors is randomly chosen for differential activation, and *all* their direct targets are assigned to the differentially transcribed gene set. Overlaying this idealized differential expression profile on the transcriptional regulatory network, we first assess how well the Fisher’s exact test applied to individual TFs recovers the original subset. For this example, we have made use of the regulatory network for *E. coli* retrieved from the RegulonDB database (Release 8.6, dated 4-11-2014), which comprises 4061 empirically validated regulatory interactions spanning 197 transcription factors and 1807 genes, covering nearly half of the coding genome [26].

We have applied one-sided Fisher’s exact test [47] to identify TFs significantly associated with the differential gene expression. This test of significance yields the probability that an overlap between the targets of a TF and the genes with altered expression at least as large as the one observed can be explained by chance. As depicted in Figure 23, the p-value calculation for each TF follows from a 2×2 contingency table that slots genes into one of four groups, which are represented by a, b, c and d. The required probability under the null hypothesis is obtained from the hypergeometric distribution, given by

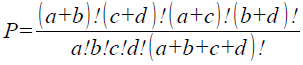

and the final p-value is a summation over terms of the above form with *a* ranging from a_min_ (the size of the overlap) to a_max_ (the size of either the differentially expressed set or the TF target set, whichever is smaller). In every trial, the over-representation p-values have been separately obtained for all regulators having non-zero overlaps with the differentially regulated gene set. The raw p-values obtained in this manner are adjusted by applying Bonferroni correction [48] to account for multiple hypotheses testing, which essentially involves multiplying the uncorrected p-values by the total number of TFs targeting at least one differentially expressed gene each. For different choices of the p-value threshold for significance, statistics for occurrences of all the TFs have been obtained over 10^4^ simulated trials.

The recovery of the input set was assessed in terms of the prediction accuracy [49], calculated as the fraction of correct classifications out of the total test set (i.e. all TFs with non-zero overlap). If *TP* denotes the number of true positives, i.e. the TFs correctly recovered from the input set, *P* stands for the total number of TFs present in the input set, *TN* is the number of TFs which are not part of the input and are not found in the enriched set, and finally *N* denotes the complement of the *P* subset, then the accuracy is given by the following formula:

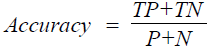

Thus, for example, if at the p < 0.05 level the enriched set of TFs is found to be identical to the input set, the accuracy assumes its maximum value of unity, and any mismatches would result in a lowered accuracy, restricted to the range 0 to 1.

### Identification of key regulators underlying differential expression by evaluating combinations of TFs

Our objective has been to identify a combination of transcription factors that best accounts for observed differential gene expression. An exhaustive search for such a combination is usually not computationally tractable. A greedy search that iteratively builds up an approximate solution based on making the best local choice at every step is routinely sought in problems of this nature. We have evaluated three variants of a heuristic procedure for obtaining an approximate solution, one of which has linear dependence of the number of p-value computations on the number of TFs in the test set, and two other alternatives which evaluate in quadratic time. These are introduced and described below.

The simplest procedure involves first ranking and ordering the TFs in accordance with their individual overlap p-values, and then, starting from the top-ranked regulator, sequentially adding the next TF from this sorted list until there is no further improvement in the combined p-value that is evaluated for the union of direct targets. We denote this heuristic as Method A, which is represented by the flowchart in Figure 4. As this scheme makes use of the original prioritized list made at the initialization to select a TF at every update, only one p-value is computed at every subsequent step, giving an O(N) dependence of the number of p-value computations on the size of the sorted list (i.e. on the number of overlapping TFs, N). Given that at each step, the best available choice is made, there is no guarantee that the converged solution represents the global minimum.

The efficacy of the above simplistic method is assessed against two other approximations, both of which involve generating a prioritized list of TFs at every step in the iteration. Thus, these approaches would have a time complexity of O(N^2^). Method B, summarized by the flowchart in Figure 5, is analogous to the greedy search that has been proposed for obtaining an approximation to the set covering problem [50]. Starting with the best single TF, as in Method A, at every subsequent step, one additional TF is added to the growing solution. In order to make this choice, we apply the criterion that, the next TF chosen is the one which has the best overlap p-value evaluated on the *remaining* subset of altered genes, i.e. those which are not already covered by the running set of TFs built up thus far. The search is stopped when no further addition of a TF from the remaining set yields a reduction in the combined p-value.

A second O(N^2^) procedure is additionally implemented. This is similar to Method B, in that a rank-ordered list of the remaining TFs is generated at every update. However, now we employ the criterion that the next TF chosen is the one which, *when added to the already built up (running) solution*, gives the best combined p-value (schematic in Figure 6). This procedure, which we shall refer to as Method C, differs not just from Method B, but also from the earlier Method A. Once convergence is reached such that no additional TF provides further improvement in the collective p-value, the progressive search is halted, yielding a candidate solution.

### Comparative analysis of approximations on published microarray data

We have performed a comparative analysis and benchmarking of the three approximations defined above on a set of microarray expression profiles downloaded from GEO [51] describing responses of wild-type *E. coli* to various stress conditions. Expression datasets associated with previous publications [52-55] for the following stress conditions for *E. coli* were downloaded: heat shock (45 °C for 10 minutes) and stationary phase; pH 5.0 and pH 8.7; sucrose osmotic stress and NaCl osmotic stress; and 1 ug/ml norfloxacin exposure, with the corresponding GEO accession numbers being GSE12190, GSE4511, GSE15534 and GSE6836 respectively. For each experiment, a set of genes differentially transcribed in a test v/s control comparison were first identified by using the accompanying GEO2R tool [56], uniformly applying an FDR threshold of <0.001 and a minimum fold-change criterion of 2.0 across all the datasets. By integrating with the RegulonDB compilation of regulatory interactions, each of the seven gene sets was then used to infer a discriminative combination of TFs by each of the three previously described methods, A, B and C.

A similar comparison among the three methods has been carried out on a larger compendium of normalized expression profiles for *E. coli* retrieved from the Many Microbes Microarrays Database (M3D, Version 4, Build 6) [57]. For the 466 experiments representing various perturbation conditions in M3D, the RMA-processed log2-transformed expression values across all the conditions were first normalized by converting them to gene-wise Z-scores, which quantify how much the expression of a gene in any particular condition deviates from the baseline defined by its mean over all the conditions. Applying a cutoff of abs(Z-score) > 2.0, for every condition, genes which show ‘abnormal’ expression with respect to their global average were identified. Thus, the approach we have followed here does not involve comparing the expression values on each array with a common control condition. Instead, the control for every gene is set by its global average.

### Ruggedness of search space and iterative search v/s steepest descent v/s stochastic search

Ruggedness of a combinatorial search space is normally defined in terms of the number of local minima accessible by steepest descent search [58-60]. Steepest gradient descent for the current problem has been implemented in the following manner: starting from a randomly selected initial configuration, at each update, the current state (TF combination) is compared with all the neighboring configurations, which each differ by the addition or exclusion of a single TF. If a neighboring configuration with a lower overlap p-value is found, then it is chosen as the new solution. This process is continued until no improvement in the p-value is obtainable by moving to a neighboring state. The probability of gradient search attaining any particular local minimum would depend on the size of the corresponding basin of attraction. In order to obtain an estimate for the ruggedness of the solution landscape, gradient descent search was run 2 × 10^3^ times starting from different initial combinations, sampled uniformly, and the number of distinct overlap p-values obtained upon convergence was tracked as a function of the number of runs carried out. This exercise was repeated for all the 7 GEO profiles introduced previously.

As an alternative approach to seeking the global minimum in a rugged search space, we have also implemented a simple formulation of simulated annealing (SA). This heuristic represents one among several different optimization techniques which contain a stochastic component allowing to overcome barriers in the search space and to avoid getting trapped in local optima [61,62]. SA modifies steepest gradient search by permitting locally non-optimal updates which can increase the p-value (the objective function here), the frequency of which is controlled by an effective temperature T. The numerical value of T modulates the probability that an update which results in a (log) p-value change of Δ(log p) is accepted, via the following formula:

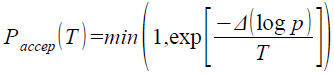

The effective temperature is gradually decreased as a function of the number of steps according to a suitably chosen annealing schedule. In the long time T → 0 limit, SA thus reduces to the usual gradient descent search. In order to circumvent issues related to optimization of the choices for temperature and annealing schedule tailored to the geometry of the search space [63], we have adopted a simplified approach by repeatedly running SA over a range of timescale settings and picking the best run in terms of the final p-value attained to represent the result of the stochastic search. The starting temperature T_i_ has been set by requiring that the initial average probability for accepting non-optimal jumps is 0.8 [61,64]. An exponential annealing schedule [65] has been adopted, parametrized by a fractional factor r which controls the rate of temperature reduction according to the formula T_n+1_ = rT_n_ at the *n*-th step. Given a final temperature T_f_(here kept fixed at 0.01 across all runs and examples), the total number of steps in one run is directly related to r by the relation N_SA_ = log_r_(T_f_/T_i_). We have repeated the search procedure for N_SA_ values ranging from 5N to 25N, and 10 runs starting from different random initial combinations were simulated for each choice of N_SA_. The running/current log p-value as well as the best p-value attained during the course of every run was recorded, and the lower of the two was chosen at the end of every run. The SA search was extended and supplemented with a gradient descent search at the end to ensure convergence to the nearest minimum in the search space. The top-ranked combination in terms of lowest log p-value obtained across the full set of runs was finally taken to represent the solution yielded by the SA approach, and used as an independent benchmark to assess the results of the previous two methods (Method C and gradient search with restarts).

### Benchmarking the proposed methodology in the idealized setting

In the artificial setting introduced earlier, we additionally run the iterative Method C on every simulated profile, to obtain a proxy for the TF combination with the least collective p-value of association. The corresponding distribution of TF occurrences across the 10^4^ trials is then compared with the distribution for the default method which was obtained earlier. The overall deviation of either distribution from the near-uniform input distribution is quantified in terms of the root mean squared error (RMSE), estimated via the formula

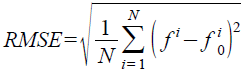

where the summation runs over all *N* TFs, and 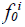 and *f^i^* stand for the proportions of trials in which the *i*-*th* TF occurs in the input and output sets respectively. A comparison between the lists of the corresponding accuracy values is also made by applying Wilcoxon signed-rank test for significant difference [66] implemented in SciPy v0.11.0.

The above exercise has been repeated, but now selecting TFs in the input set to be consistent with the regulatory network structure of TF-TF regulatory interactions. In other words, we start with a ‘seed’ set composed of a small number of randomly selected TFs, and then add all the other TFs in the network which lie downstream of TFs in the seed set and are therefore regulated directly or indirectly by them. This is done under the assumption that perturbing the activity of one TF is expected to lead to a cascade of regulatory rewiring downstream over a longer timescale (in the previous analysis, the effect of perturbing TFs was assumed to be limited to only the direct downstream targets). As before, 10^4^ seed sets were randomly sampled.

In order to confirm whether the trends are sensitive to the presence of misclassification (noise) distorting the expression pattern, we have revised the earlier deterministic setting by now adding noise in the form of misclassification of genes. This has been implemented by randomly selecting a fraction of the genes in the network and reassigning them to the opposite class. Thus, if a differentially expressed gene is chosen, it is reclassified as unaltered, and vice versa. The performance of the two inference methods (default and Method C) was assessed over 10^4^ random trials in the presence of 5% misclassification rate.

### Independent comparative assessment on *M. tuberculosis* TF over-expression experimental data

As a final and independent assessment of the presented methodology on a different organism, we have applied it vis-à-vis the standard approach in the context of *M. tuberculosis* (MTB) gene expression data. Microarray profiles made available as part of a recently published large-scale study [67] were downloaded from [68]. This dataset comprises over-expression phenotypes spanning 206 MTB TFs, and genes showing an up/down regulation log fold change greater than 1.0 were called differentially expressed in each experiment. This expression change information was integrated with a curated transcriptional regulatory network composed of 91 TFs and 3682 regulatory connections covering 1787 MTB genes in total. This reference network has been obtained by merging two independent previously published studies [69,70], and was assembled by a combination of curated experimental evidences and orthology assignment from related species (*E. coli* and *C. glutamicum*). As in the previous analyses, the standard testing of individual TFs as well as the Method C were applied to each gene list, and recovery performance was assessed in terms of the number of times the causal TF (known *a priori*) is statistically enriched at p < 0.05 confidence threshold, relative to the number of times it is present in the TF combination yielded by the iterative search.

**Figure 1.**
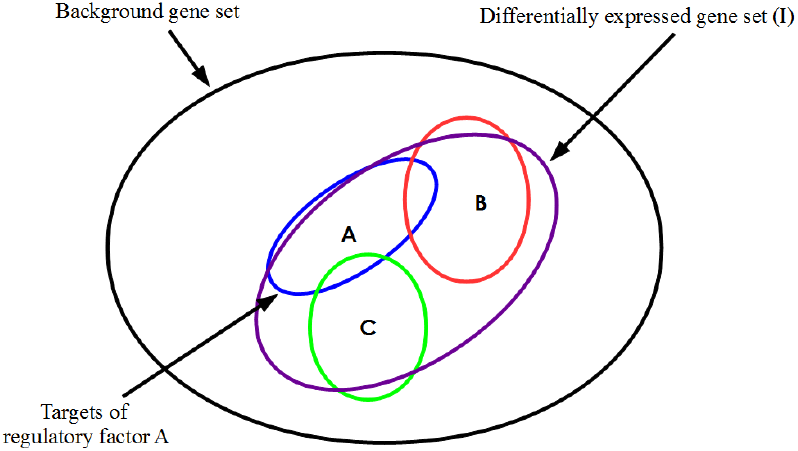
Differential expression profile overlaid on a regulatory network. Venn diagram illustrating an example situation involving three transcription factors A, B and C each of which targets a distinct group of genes. Each target set also has an overlap with the input set I, which represents the genes detected as differentially expressed in a case v/s control comparison of transcriptomic data.

**Figure 2.**
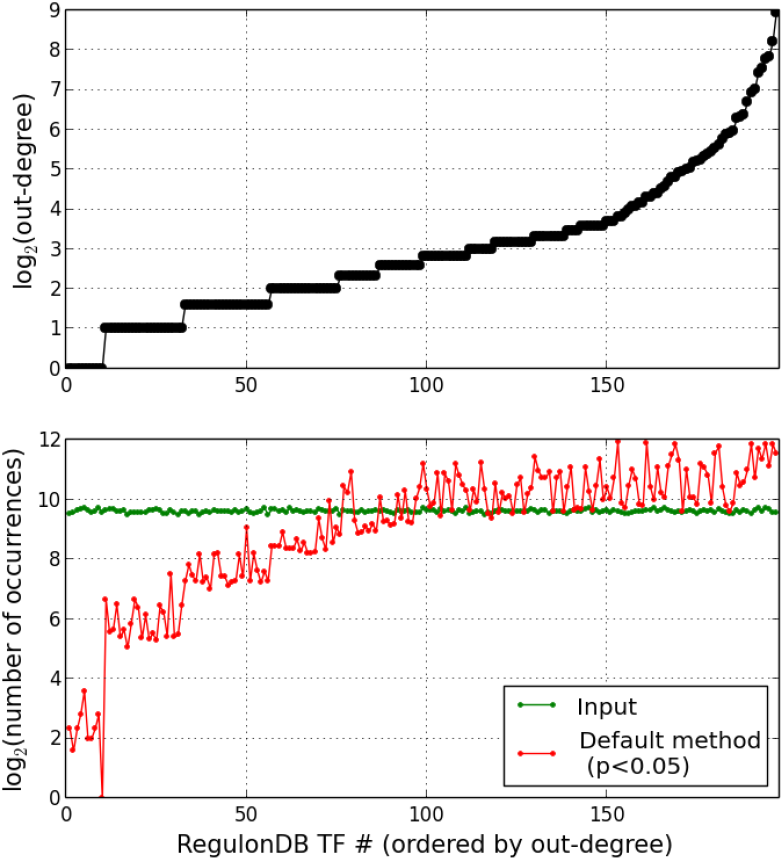
Transcription factor enrichment from simulated trials. (A) The *E. coli* regulatory meta-network used contains 197 TFs, exhibiting a broad out-degree distribution. (B) Distribution of transcription factor occurrences in the input and statistically enriched sets, obtained by aggregating 10^4^ randomly generated binarized profiles in the noiseless limit.

**Figure 3.**
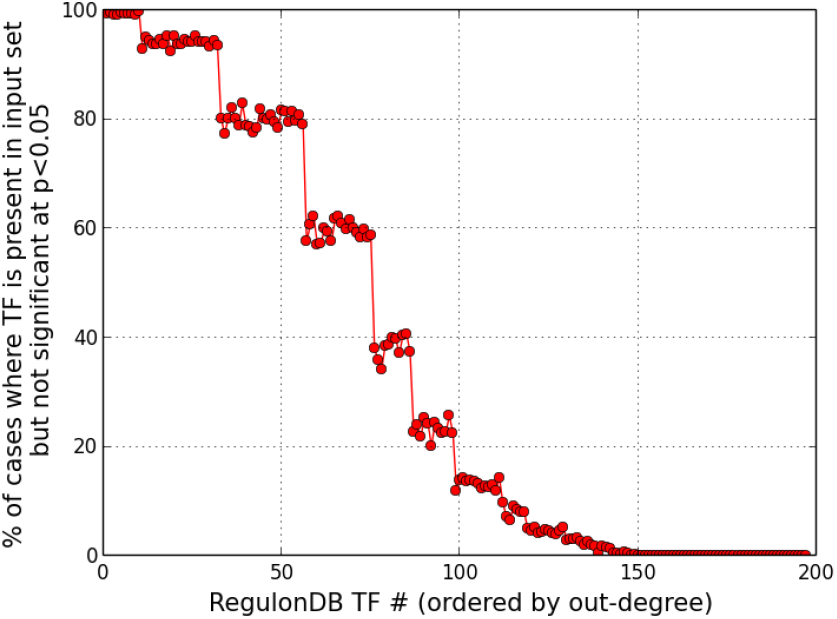
Under-representation of TFs in the enriched set. TFs with smaller target sets may not always be statistically enriched, if multiple TFs are simultaneously ‘perturbed’ producing an extensively altered expression profile.

**Figure 4.**
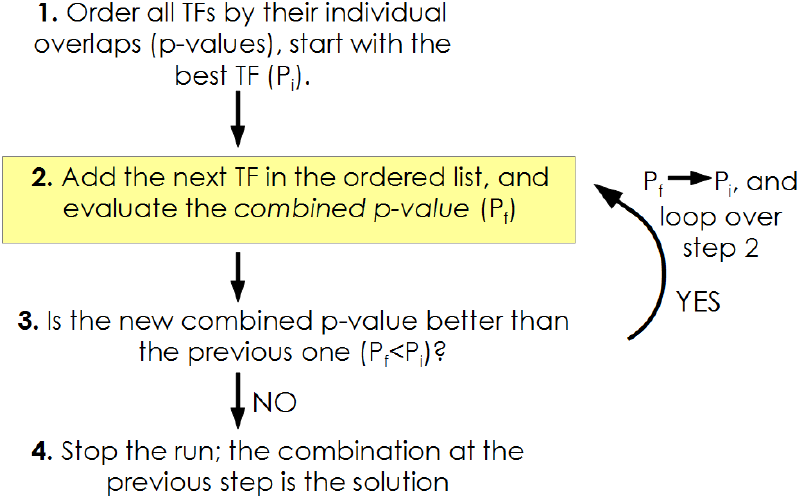
Schematic representation of the linear (O(N)) procedure, Method A, for obtaining an approximation for the most predictive subnetwork.

**Figure 5.**
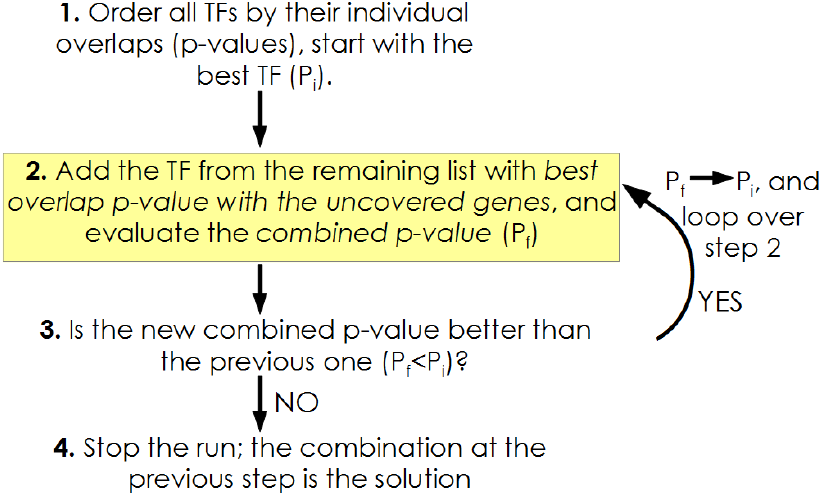
Schematic of the heuristic adapted from the set cover problem, Method B, with a running time (number of p-value evaluations) quadratic in the number of TFs.

**Figure 6.**
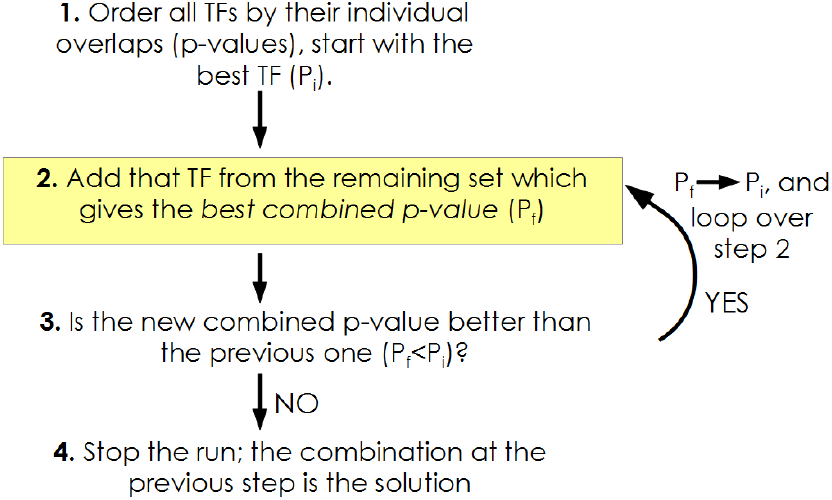
Flowchart for the approximate Method C, which also involves O(N^2^) p-value computations.

**Figure 7.**
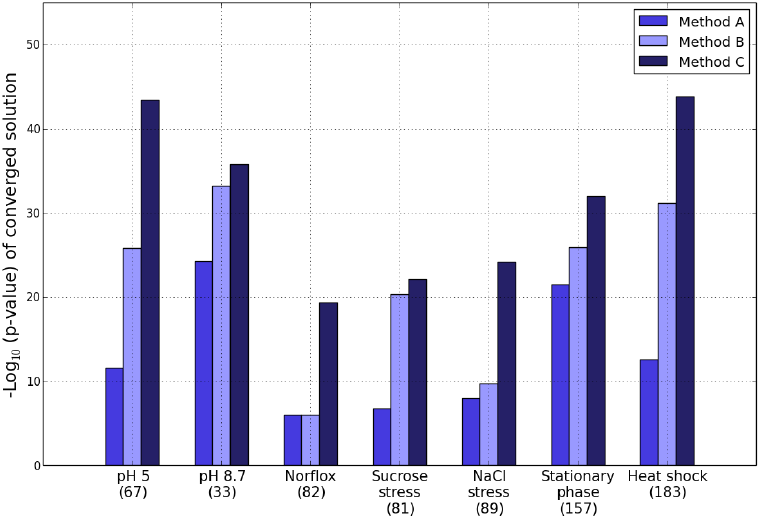
Benchmarking the iterative methods on GEO microarray datasets. Comparison of the iterative methods A-C to identify the maximally discriminative combination of TFs, applied to 7 *E. coli* expression datasets retrieved from GEO. The number in brackets under every label corresponds to the number of TFs in the background network targeting at least one differentially expressed gene each.

**Figure 8.**
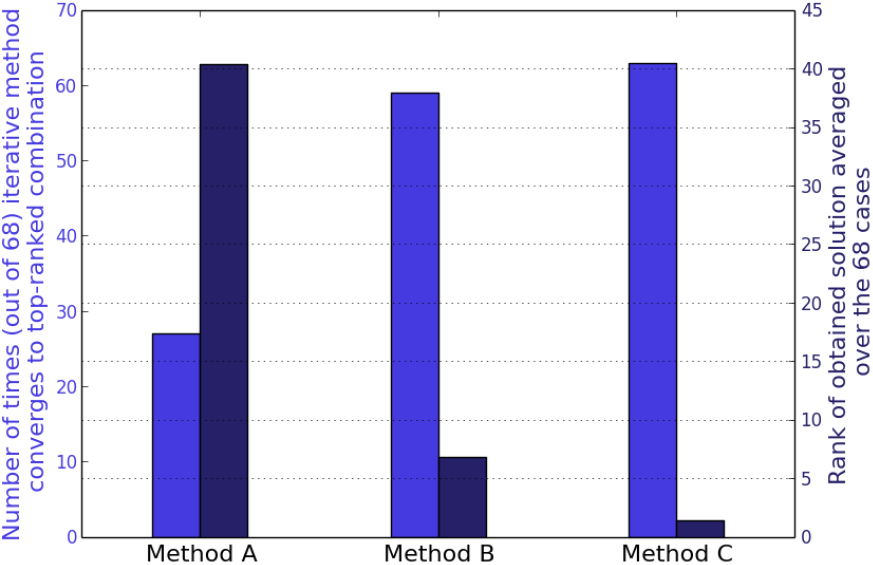
Benchmarking the iterative methods on M3D data. Comparison of the three heuristics applied to a subset of M3D expression profiles where the manageable numbers of overlapping TFs make exhaustive enumeration possible (Z-score threshold for significance of altered expression = ± 2.0).

**Figure 9.**
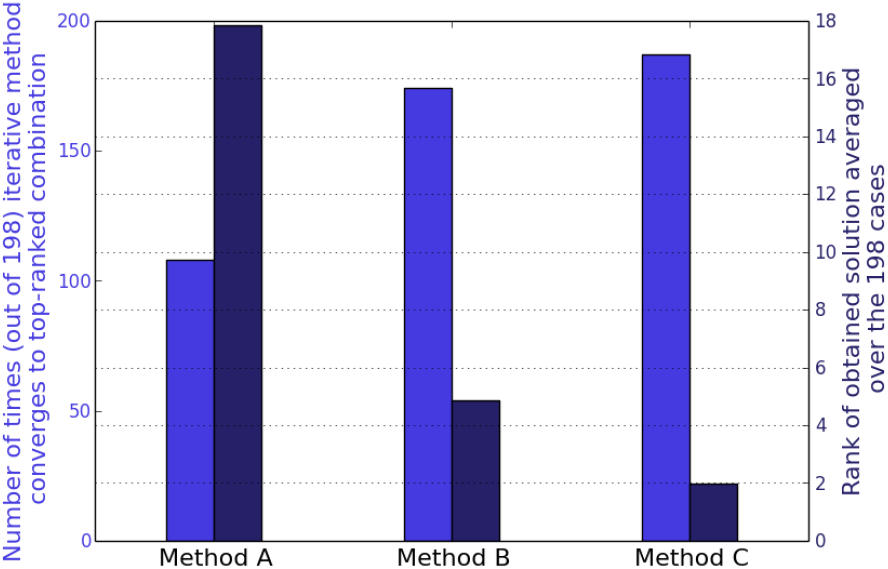
Benchmarking the iterative methods on M3D data. Same comparison as in Fig. 8, but now with application of a more stringent Z-score significance threshold of ± 3.0. 198 conditions were identified as having not more than 14 overlapping TFs, making it straightforward to carry out exhaustive search.

**Figure 10.**
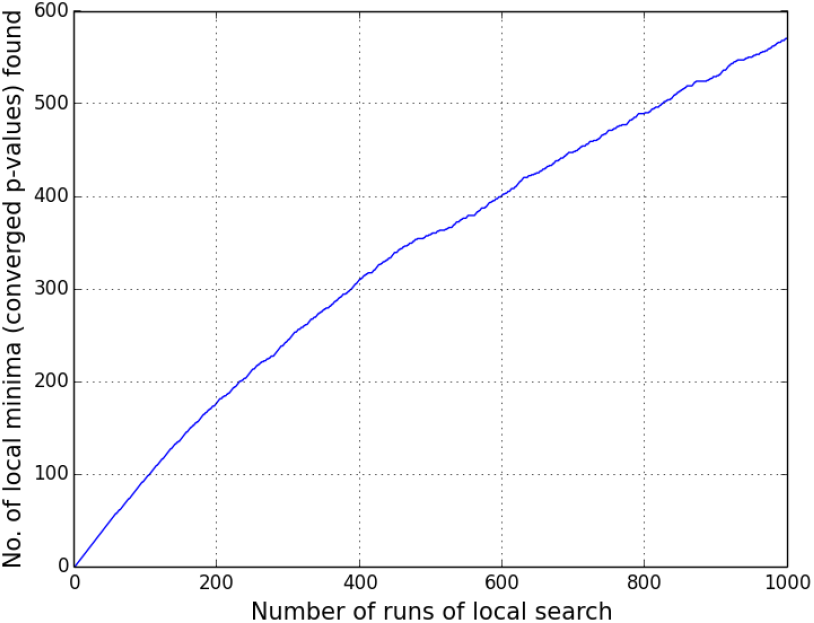
Ruggedness of problem search space. Ruggedness estimated by the number of local minima attained by steepest descent search as a function of the number of initial conditions sampled. This figure is for the heat shock response profile.

**Figure 11.**
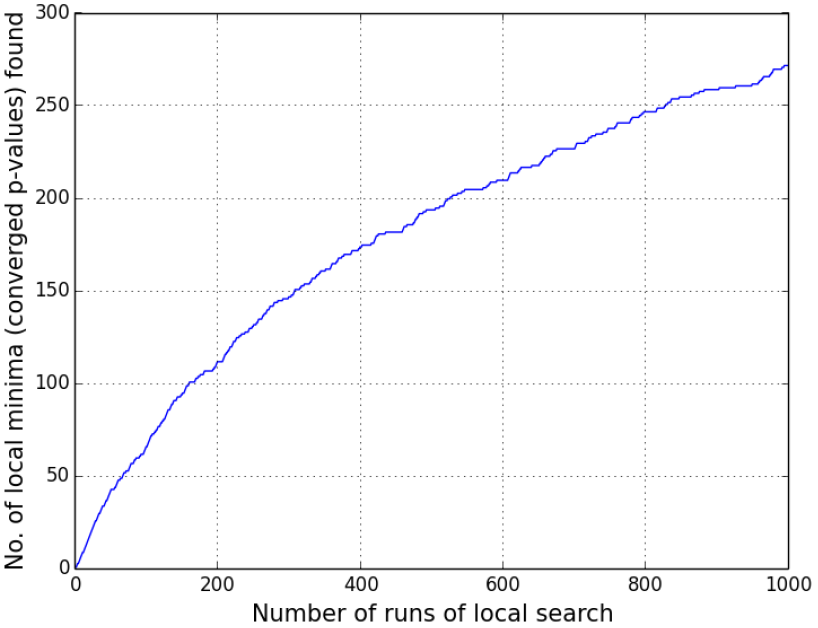
Sampling of local minima by gradient descent search for the stationary phase comparison dataset.

**Figure 12.**
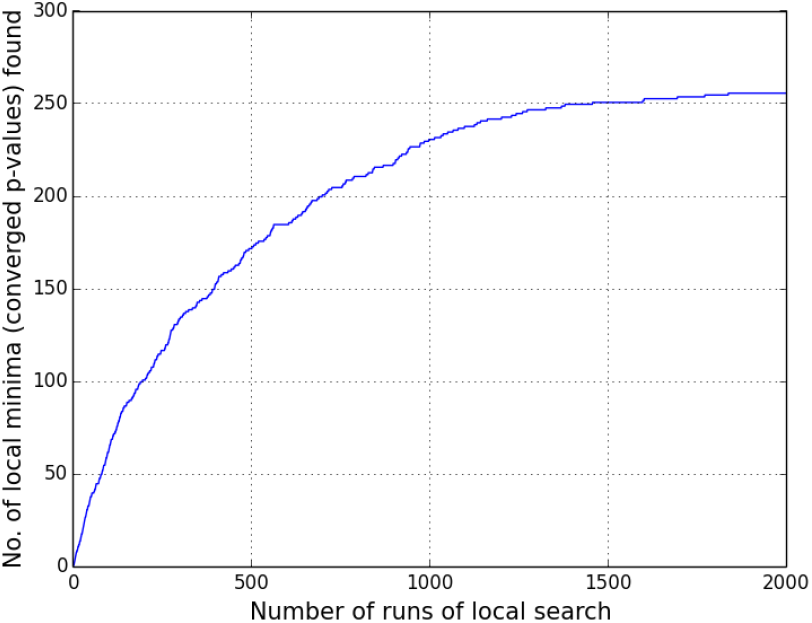
Sampling of local minima by gradient descent search for the low pH (5.0) comparison dataset.

**Figure 13.**
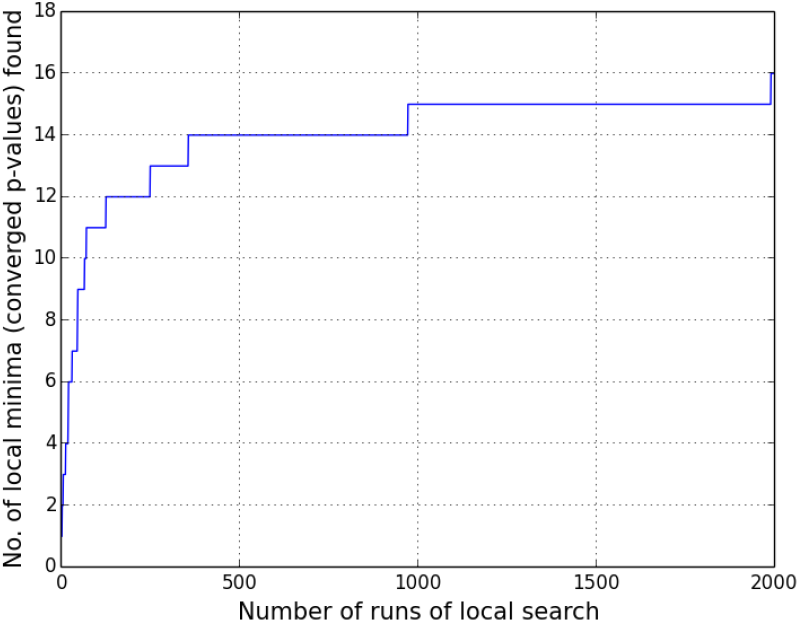
Sampling of local minima by gradient descent search for the high pH (8.7) comparison dataset.

**Figure 14.**
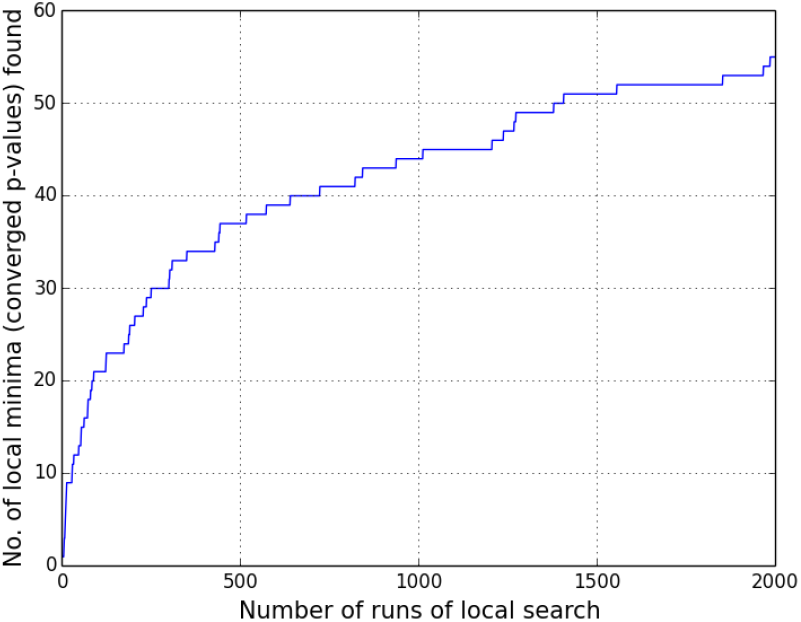
Sampling of local minima by gradient descent search for the norfloxacin exposure dataset.

**Figure 15.**
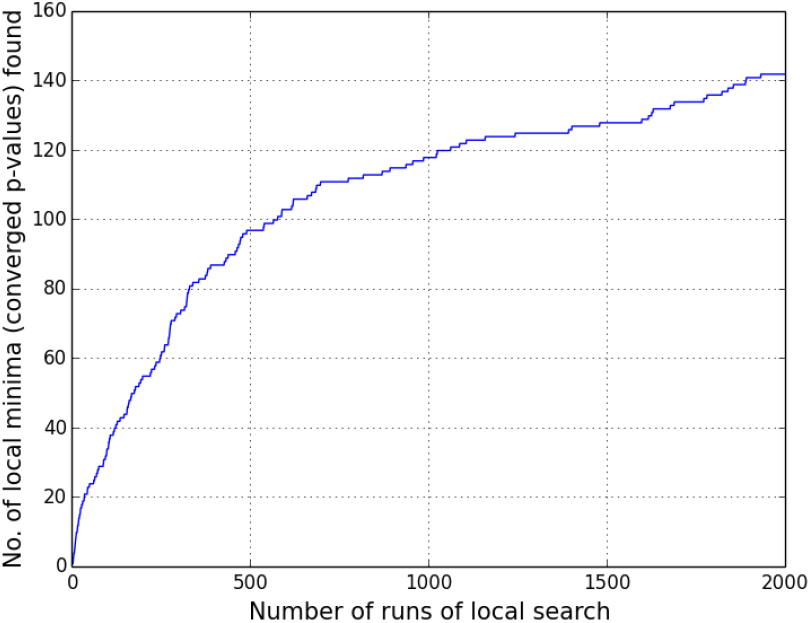
Sampling of local minima by gradient descent search for the NaCl osmotic stress response dataset.

**Figure 16.**
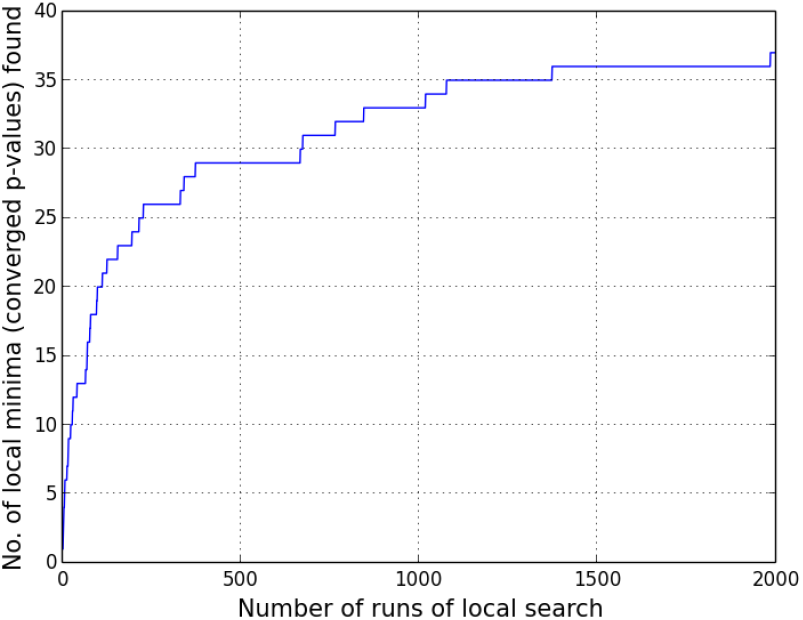
Sampling of local minima by gradient descent search for the sucrose osmotic stress response dataset.

**Figure 17.**
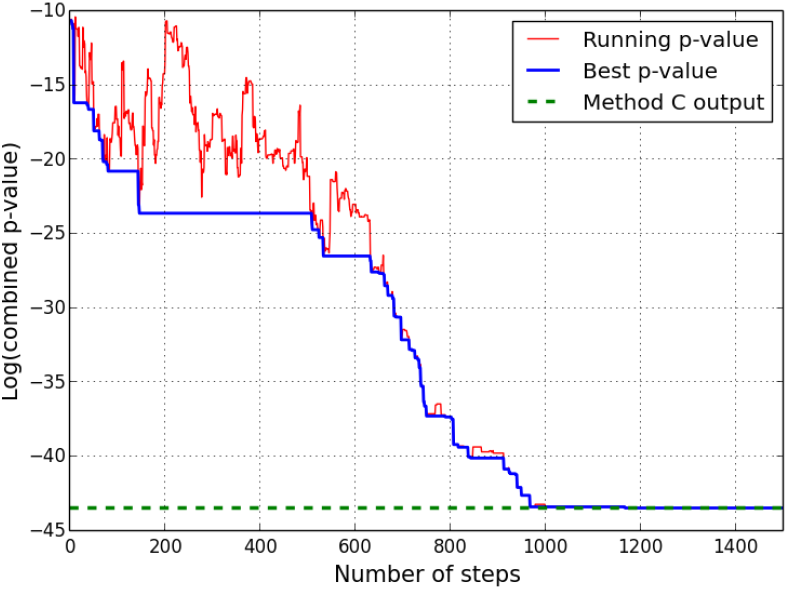
Example of simulated annealing search. A single run of SA stochastic search for the global minimum p-value, applied to the pH = 5 vs. pH =7 comparison microarray dataset. Sharp fluctuations in the red curve arise from the possibility of non-optimal updates, the propensity for which depends on the (gradually decreasing) annealing temperature.

**Figure 18.**
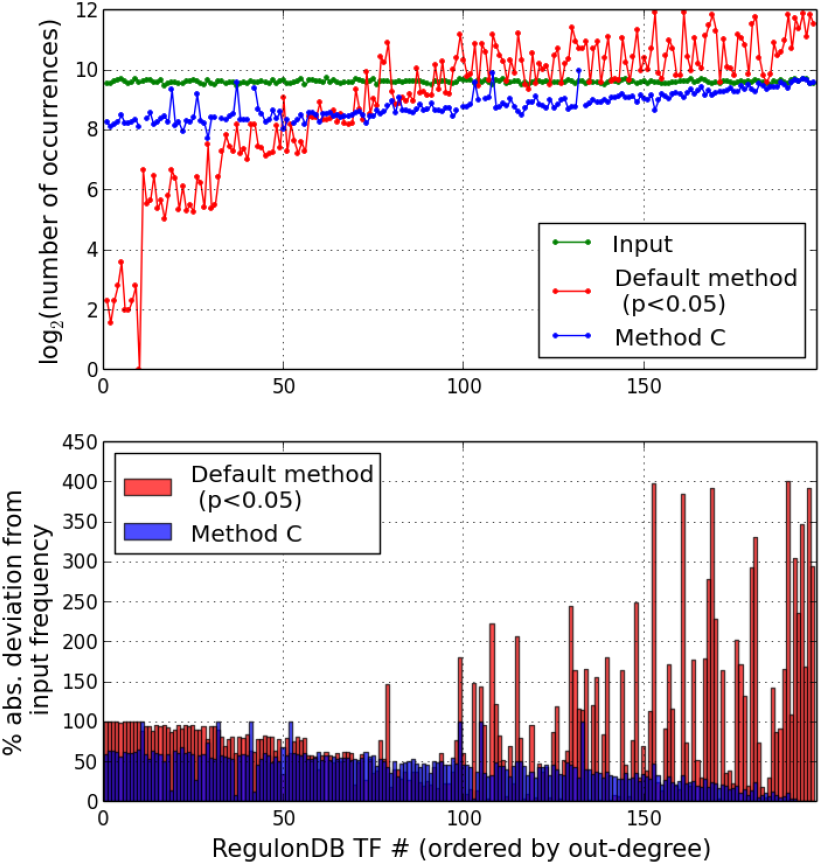
Comparison of the combinatorial approach with the default method on simulated trials. (A) Comparison of default method with the iterative Method C (over 10^4^ trials) in the noiseless limit which was introduced in Fig. 2. (B) Deviations of the red and blue curves from the green profile in (A) have been represented in terms of profiles of percentage differences.

**Figure 19.**
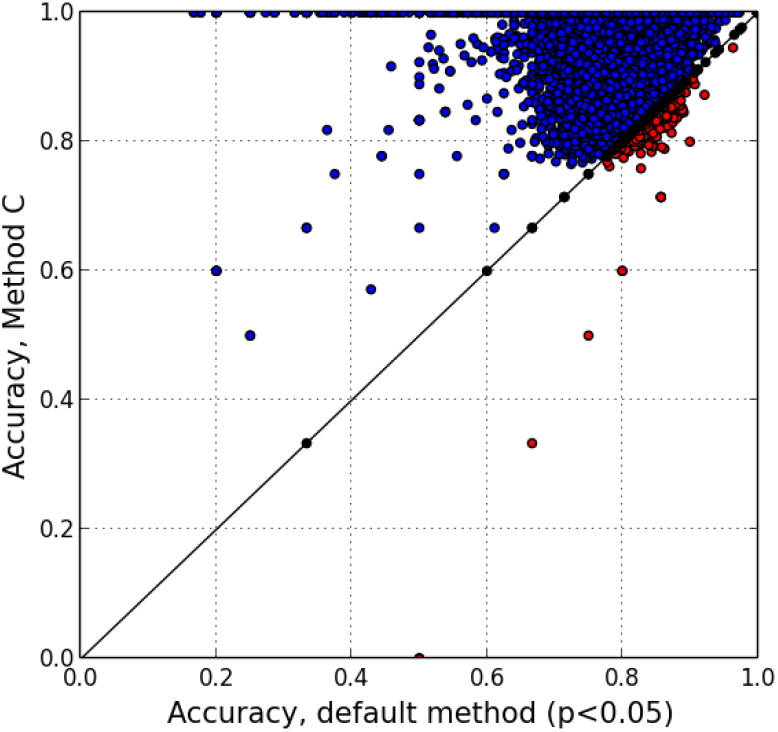
Recovery accuracy for TFs based on simulated trials. Comparison of classification accuracy values yielded by the default and iterative approaches across the 10^4^ random trials, represented as a two-dimensional scatter plot. Points above and below the equality line are colored blue and red respectively, while those lying on the 45° diagonal are shownin black.

**Figure 20.**
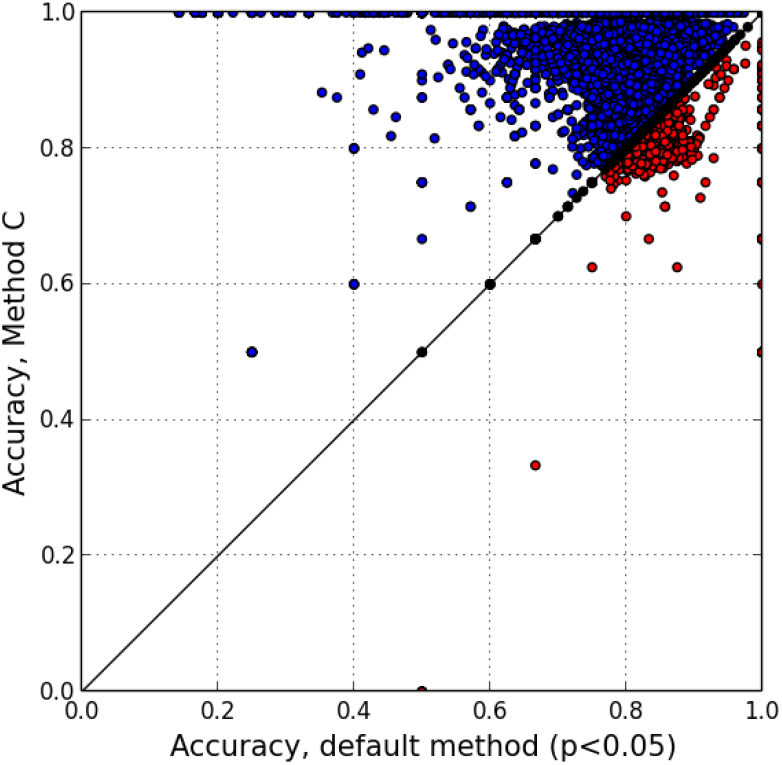
TF recovery accuracy based on simulated trials. Paired comparison of accuracy values yielded by the default and combinatorial approaches across 10^4^ random trials in which the input set of TFs is selected in such a way as to be consistent with the underlying hierarchy of TF-TF regulatory interactions in the RegulonDB network.

**Figure 21.**
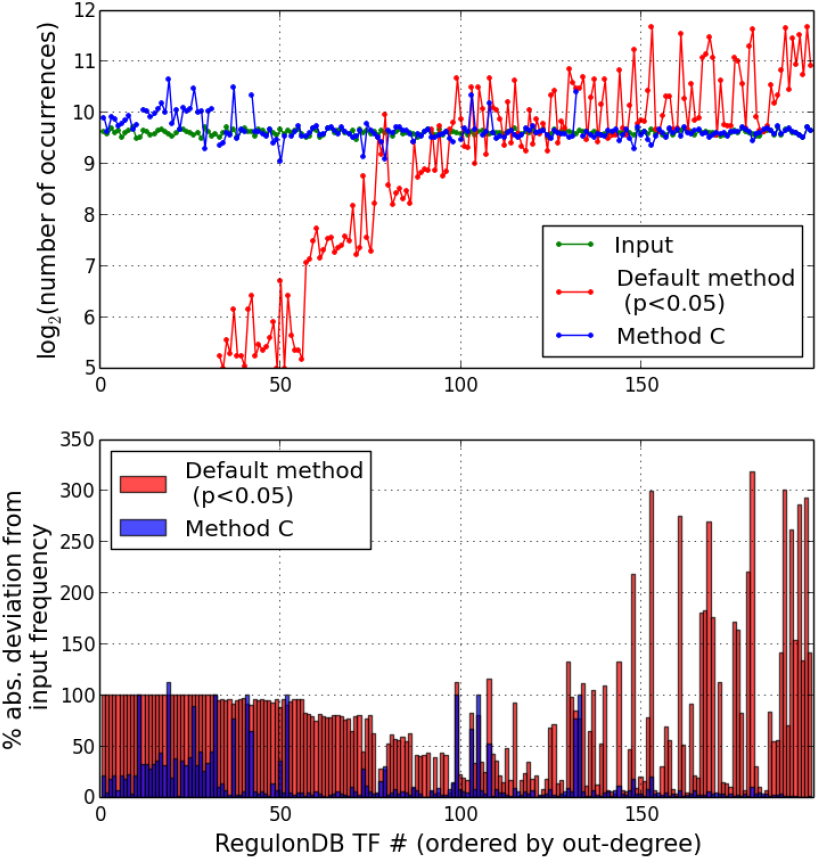
Comparison of the combinatorial approach with the default method on simulated trials with inclusion of noise. Distribution of TF occurrences for the case where the idealized binary differential expression profile is distorted by addition of 5% misclassification in gene assignment. Averaged over 10^4^ simulated trials.

**Figure 22.**
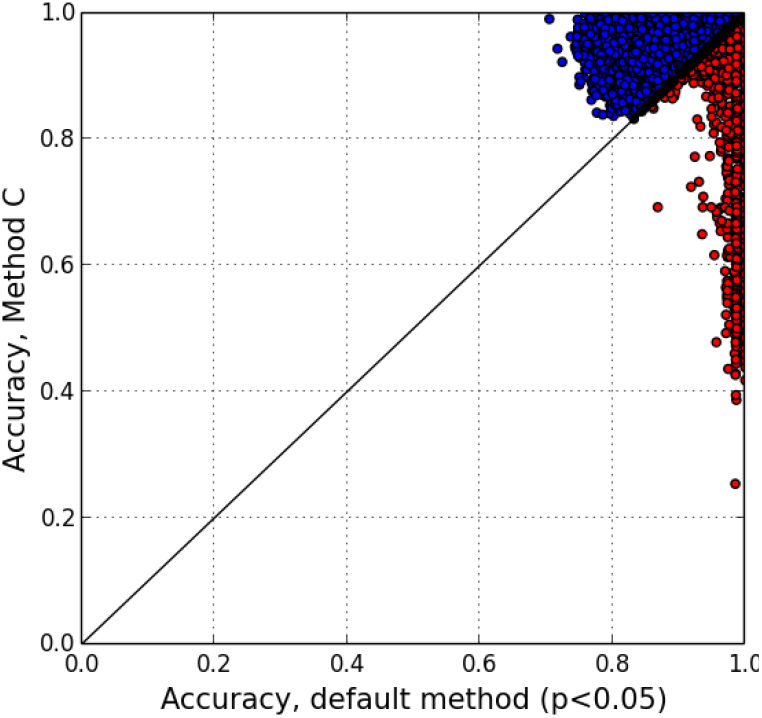
TF recovery accuracy based on simulated trials with noise. Paired comparison of accuracy values across trials where 5% classification error is added to the binarized differential expression profile.

**Figure 23.**
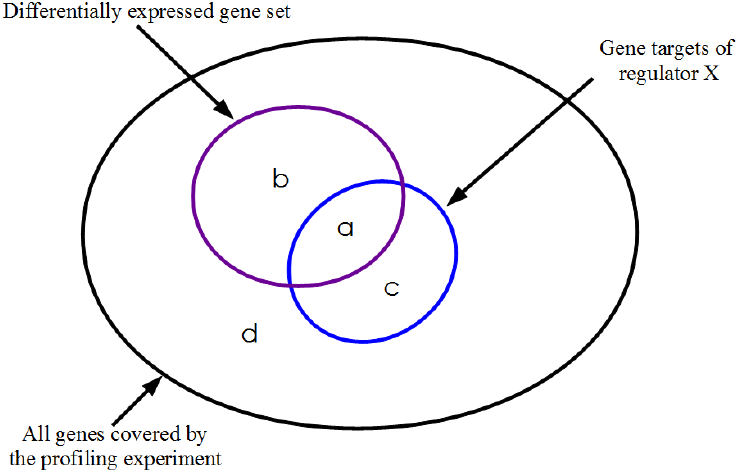
Transcription factor enrichment from large scale differential expression data. Graphical representation of the entries a, b, c and d in the 2×2 contingency table on which the computation of the enrichment p-value via Fisher’s exact test is based. This test evaluates the probability that the size of the observed overlap (a) is consistent with random distribution of the differentially expressed subset.

## Acknowledgements

Financial support from the Department of Biotechnology, Government of India (Grant SM/BT/01/07/02/2008) and Department of Science and Technology, Government of India (Grant INT/RUS/RFBR/P-154) is gratefully acknowledged.

## REFERENCES

1. Soon WW, Hariharan M, Snyder MP. High-throughput sequencing for biology and medicine. Mol. Syst. Biol. 2013, 9:640.

2. Vidal M, Cusick ME, Barabási AL. Interactome networks and human disease. Cell 2011, 144(6):986–998.

3. Barabási A-L, Oltvai ZN. Network biology: understanding the cell’s functional organization. Nature Rev. Genet. 2004, 5(2):101–113.

5. Wu C, Zhu J, Zhang X. Integrating gene expression and protein-protein interaction network to prioritize cancer-associated genes. BMC Bioinformatics 2012, 13:182.

6. Functional Genomics. In: Nature insight. Nature 2000, 405:819–85.

7. Ball CA, Brazma A, Causton H, Chervitz S, Edgar R, et al. Submission of microarray data to public repositories. PLoS Biol 2004, 2(9): E317.

8. Metzker ML. Sequencing technologies - the next generation. Nat Rev Genet 2010, 11(1):31–46.

9. Alon U. An introduction to systems biology: design principles of biological circuits. Boca Raton: Chapman & Hall/CRC; 2007.

10. Kim HD, Shay T, O’Shea, EK, Regev A. Transcriptional regulatory circuits: predicting numbers from alphabets. Science 2009, 325(5939):429–432.

11. Blais A, Dynlacht BD. Constructing transcriptional regulatory networks. Genes Dev. 2005, 19(13): 1499–1511.

12. ENCODE Project Consortium, Birney E, Stamatoyannopoulos JA, Dutta A, Guigó R, et al. Identification and analysis of functional elements in 1% of the human genome by the ENCODE pilot project. Nature 2007, 447(7146):799–816.

13. Badis G, Berger MF, Philippakis AA, Talukder S, Gehrke AR, et al. Diversity and complexity in DNA recognition by transcription factors. Science 2009, 324(5935):1720–1723.

14. Stormo GD. DNA binding sites: representation and discovery. Bioinformatics 2000, 16(1): 16–23.

15. D'haeseleer P. How does DNA sequence motif discovery work? Nat Biotechnol 2006, 24(8):959–961.

16. Kharchenko PV, Tolstorukov MY, Park PJ. Design and analysis of ChIP-seq experiments for DNA-binding proteins. Nature Biotechnol 2008, 26(12): 1351–1359.

17. Daily K, Patel VR, Rigor P, Xie X, Baldi P. MotifMap: integrative genome-wide maps of regulatory motif sites for model species. BMC Bioinformatics 2011, 12:495.

18. Lee TI, Rinaldi NJ, Robert F, Odom DT, Bar-Joseph Z, et al. Transcriptional regulatory networks in Saccharomyces cerevisiae. Science 2002, 298(5594):799–804.

19. Teixeira MC, Monteiro P, Jain P, Tenreiro S, Fernandes AR, et al. The YEASTRACT database: a tool for the analysis of transcription regulatory associations in Saccharomyces cerevisiae. Nucleic Acids Res. 2006, 34(Database issue):D446–D451.

20. Murali T, Pacifico S, Yu J, Guest S, Roberts GG, Finley RL. DroID 2011: a comprehensive, integrated resource for protein, transcription factor, RNA and gene interactions for Drosophila. Nucleic Acids Res. 2011, 39(Database issue):D736–D743.

21. ENCODE Project Consortium, Dunham I, Kundaje A, Aldred SF, Collins PJ, et al. An integrated encyclopedia of DNA elements in the human genome. Nature 2012, 489(7414):57–74.

22. Mouse ENCODE Consortium, Stamatoyannopoulos JA, Snyder M, Hardison R, Ren B, et al. An encyclopedia of mouse DNA elements (Mouse ENCODE). Genome Biol 2012, 13(8):418.

23. Wang J, Zhuang J, Iyer S, Lin X, Whitfield TW, et al. Sequence features and chromatin structure around the genomic regions bound by 119 human transcription factors. Genome Res. 2012, 22(9):1798–1812.

24. Bovolenta LA, Acencio ML, Lemke N. HTRIdb: an open-access database for experimentally verified human transcriptional regulation interactions. BMC Genomics 2012, 13:405.

25. Liberzon A, Subramanian A, Pinchback R, Thorvaldsdottir H, Tamayo P, Mesirov JP. Molecular signatures database (MSigDB) 3.0. Bioinformatics 2011, 27(12):1739–1740.

26. Salgado H, Peralta-Gil M, Gama-Castro S, Santos-Zavaleta A, Muñiz-Rascado L, et al. RegulonDB v8.0: Omics data sets, evolutionary conservation, regulatory phrases, cross-validated gold standards and more. Nucleic Acids Res. 2013, 41(Database issue):D203–D213.

27. Wingender E, Chen X, Hehl R, Karas H, Liebich I, et al. TRANSFAC: an integrated system for gene expression regulation. Nucleic Acids Res. 2000, 28(1):316–319.

28. Matys V, Kel-Margoulis OV, Fricke E, Liebich I, Land S, et al. TRANSFAC and its module TRANSCompel: transcriptional gene regulation in eukaryotes. Nucleic Acids Res. 2006, 34(Database issue):D108–D110.

29. Mathelier A, Zhao X, Zhang AW, Parcy F, Worsley-Hunt R, et al. JASPAR 2014: an extensively expanded and updated open access database of transcription factor binding profiles. Nucleic Acids Res. 2014, 42 (Database issue):D142–D147.

30. Kulakovskiy IV, Medvedeva YA, Schaefer U, Kasianov AS, Vorontsov IE, et al. HOCOMOCO: a comprehensive collection of human transcription factor binding sites models. Nucleic Acids Res. 2013, 41(Database issue):D195–D202.

31. Hume MA, Barrera LA, Gisselbrecht SS, Bulyk ML. UniPROBE, update 2015: new tools and content for the online database of protein-binding microarray data on protein-DNA interactions. Nucleic Acids Res. 2014. doi: 10.1093/nar/gku1045.

32. Lachmann A, Xu H, Krishnan J, Berger SI, Mazloom AR, Ma'ayan A. ChEA: transcription factor regulation inferred from integrating genome-wide ChIP-X experiments. Bioinformatics 2010, 26(19):2438–44.

33. Kou Y, Chen EY, Clark NR, Tan CM, Ma‘ayan A. ChEA2: Gene-Set Libraries from ChIP-X Experiments to Decode the Transcription Regulome. Multidisciplinary Research and Practice for Information Systems. CD-ARES 2013. Lecture Notes in Computer Science 2013, 8127:416–430.

34. Lachmann A, Ma'ayan A. KEA: Kinase Enrichment Analysis. Bioinformatics 2009, 25(5):684–6.

35. Chen EY, Xu H, Gordonov S, Lim MP, Perkins MH, Ma'ayan A. Expression2Kinases: mRNA Profiling Linked to Multiple Upstream Regulatory Layers. Bioinformatics 2012, 28(1): 105–111.

36. Auerbach RK, Chen B, Butte AJ. Relating genes to function: identifying enriched transcription factors using the ENCODE ChIP-Seq significance tool. Bioinformatics 2013, 29(15):1922–1924.

37. Bleda M, Medina I, Alonso R, De Maria A, Salavert F, Dopazo J. Inferring the regulatory network behind a gene expression experiment. Nucleic Acids Res. 2012, 40(Web Server issue):W168–W172.

38. Zhang B, Kirov SA, Snoddy JR. WebGestalt: an integrated system for exploring gene sets in various biological contexts. Nucleic Acids Res. 2005, 33(Web Server issue):W741–748.

39. Dubchak I, Munoz M, Poliakov A, Salomonis N, Minovitsky S, Bodmer R, Zambon AC. Whole-Genome rVISTA: a tool to determine enrichment of transcription factor binding sites in gene promoters from transcriptomic data. Bioinformatics 2013, 29(16):2059–61.

40. Loots GG, Ovcharenko I, Pachter L, Dubchak I, Rubin EM. rVista for comparative sequence-based discovery of functional transcription factor binding sites. Genome Res. 2002, 12(5):832–839.

41. Elkon, R, Linhart C, Sharan R, Shamir R, Shiloh Y. Genome-wide In-silico Identification of Transcriptional Regulators Controlling Cell Cycle in Human Cells. Genome Res. 2003, 13(5):773–780.

42. Frith MC, Fu Y, Yu L, Chen J-F, Hansen U, Weng Z. Detection of functional DNA motifs via statistical over-representation. Nucleic Acids Res. 2004, 32(4):1372–81.

43. Haverty PM, Hansen U, Weng Z. Computational inference of transcriptional regulatory networks from expression profiling and transcription factor binding site identification. Nucleic Acids Res. 2004, 32(1):179–188.

44. Smyth G. Linear models and empirical bayes methods for assessing differential expression in microarray experiments. Stat. Appl. Genet. Mol. Biol. 2004, 3:3.

45. Tusher, VG, Tibshirani R, et al. Significance analysis of microarrays applied to the ionizing radiation response. Proc. Natl. Acad. Sci. U S A 2001, 98(9):5116–5121.

46. Jeffery IB, Higgins DG, Culhane AC. Comparison and evaluation of methods for generating differentially expressed gene lists from microarray data. BMC Bioinformatics 2006, 7:359.

47. Fisher RA. On the interpretation of x^2^ from contingency tables, and the calculation of P. J. Royal Stat. Soc. 1922, 85(1):87–94.

48. Bland JM, Altman DG. Multiple significance tests: The Bonferroni method. BMJ 1995, 310 (6973):170.

49. Tarca AL, Carey VJ, Chen XW, Romero R, Drăghici S. Machine Learning and Its Applications to Biology. PLoS Comput. Biol. 2007, 3(6):e116.

50. Chvatal VA. Greedy Heuristic for the Set-Covering Problem. Mathematics of Operations Research 1979, 4(3):233–235.

51. Edgar R, Domrachev M, Lash AE. Gene Expression Omnibus: NCBI gene expression and hybridization array data repository. Nucleic Acids Res. 2002, 30(10):207–210.

52. Cho BK, Zengler K, Qiu Y, Park YS, Knight EM, et al. The transcription unit architecture of the Escherichia coli genome. Nucleic Acids Res. 2009, 27(11):1043–1049.

53. Maurer LM, Yohannes E, Bondurant SS, Radmacher M, Slonczewski JL. pH regulates genes for flagellar motility, catabolism, and oxidative stress in Escherichia coli K-12. J. Bacteriol. 2005, 187(1):304–319.

54. Shabala L, Bowman J, Brown J, Ross T, McMeekin T, Shabala S. Ion transport and osmotic adjustment in Escherichia coli in response to ionic and non-ionic osmotica. Environ. Microbiol. 2009, 11(1): 137–148.

55. Faith JJ, Hayete B, Thaden JT, Mogno I, Wierzbowski J, et al. Large-scale mapping and validation of Escherichia coli transcriptional regulation from a compendium of expression profiles. PLoS Biol. 2007, 5(1):e8.

56. Barrett T, Wilhite SE, Ledoux P, Evangelista C, Kim IF, et al. NCBI GEO: archive for functional genomics data sets—update. Nucleic Acids Res. 2013, 41(Database issue):D991–D995.

57. Faith JJ, Driscoll ME, Fusaro VA, Cosgrove EJ, Hayete B, et al. Many Microbe Microarrays Database: uniformly normalized Affymetrix compendia with structured experimental metadata. Nucleic Acids Res. 2008, 36(Database issue):D866–70.

58. Kouyos RD, Leventhal GE, Hinkley T, Haddad M, Whitcomb JM, et al. Exploring the Complexity of the HIV-1 Fitness Landscape. PLoS Genet. 2012, 8(3):e1002551.

59. Arjan J, De Visser GM, Krug J. Empirical fitness landscapes and the predictability of evolution. Nature Rev. Genet. 2014, 15(7):480–490.

60. Kauffman S, Levin S. Towards a general theory of adaptive walks on rugged landscapes. J Theor Biol 1987, 128(1):11–45.

61. Kirkpatrick S, Gerlatt Jr CD, Vecchi MP. Optimization by Simulated Annealing. IBM Research Report RC 9355, 1982.

62. Weisbuch G. Complex Systems Dynamics. Santa-Fe Institute Studies in the Sciences of Complexity, Addison-Wesley, Redwood City, CA, USA; 1990.

63. Bounds DG. New Optimization Methods from Physics and Biology. Nature 1987, 329:215–218.

64. Kirkpatrick S. Optimization by Simulated Annealing: Quantitative Studies. J. Stat. Phys. 1984, 34(5-6):975–986.

65. Kirkpatrick S, Gelatt CD Jr, Vecchi MP. Optimization by Simulated Annealing. Science 1983, 220(4598):671–680.

66. Wilcoxon F. Individual comparisons by ranking methods. Biometrics Bulletin 1945, 1(6):80–83.

67. Rustad TR, Minch KJ, Ma S, Winkler JK, Hobbs S, et al. Mapping and manipulating the Mycobacterium tuberculosis transcriptome using a transcription factor overexpression-derived regulatory network. Genome Biol. 2014, 15(11):502.

68. http://networks.systemsbiology.net/mtb/data-center.

69. Rohde KH, Veiga DF, Caldwell S, Balázsi G, Russell DG. Linking the Transcriptional Profiles and the Physiological States of Mycobacterium tuberculosis during an Extended Intracellular Infection. PLoS Pathog. 2012, 8(6):e1002769.

70. Sanz J, Navarro J, Arbués A, Martín C, Marijuán PC, et al. The Transcriptional Regulatory Network of Mycobacterium tuberculosis. PLoS ONE 2011, 6(7):e22178.

71. Huang da W, Sherman BT, Lempicki RA. Bioinformatics enrichment tools: paths toward the comprehensive functional analysis of large gene lists. Nucleic Acids Res. 2009, 37(1):1–13.

72. Dennis G Jr, Sherman BT, Hosack DA, Yang J, Gao W, et al. DAVID: database for annotation, visualization, and integrated discovery. Genome Biol. 2003, 4(5):P3.

73. Ashburner M, Ball CA, Blake JA, Botstein D, Butler H, et al. Gene Ontology: tool for the unification of biology. Nature Genet. 2000, 25(1):25–29.

74. Reimand J, Kull M, Peterson H, Hansen J, Vilo J. g:Profiler -- a web-based toolset for functional profiling of gene lists from large-scale experiments. Nucleic Acids Res. 2007, 35(Web Server issue):W193–W200.

75. Subramanian A, Tamayo P, Mootha VK, Mukherjee S, Ebert BL, et al. Gene set enrichment analysis: a knowledge-based approach for interpreting genome-wide expression profiles. Proc. Natl. Acad. Sci. U S A 2005, 102(43):15545–15550.

76. Irizarry RA, Wang C, Zhou Y, Speed TP. Gene Set Enrichment Analysis Made Simple. Stat. Methods Med. Res. 2009, 18(6):565–575.

77. Lee HK, Braynen W, Keshav K, Pavlidis P. ErmineJ: tool for functional analysis of gene expression data sets. BMC Bioinformatics 2005, 6:269.

78. Chuang HY, Lee E, Liu YT, Lee D, Ideker T. Network-based classification of breast cancer metastasis. Mol. Syst. Biol. 2007, 3:140.

79. Box GEP, Jenkins GM, Reinsel GC. Time Series Analysis: Forecasting and Control. 3rd ed. Englewood Cliffs, NJ, Prentice Hall 1994.

80. Balaji S, Babu MM, Iyer LM, Luscombe NM, Aravind L. Comprehensive analysis of combinatorial regulation using the transcriptional regulatory network of yeast. J. Mol. Biol. 2006, 360(1):213–227.

81. Kim J, Choi M, Kim J-R, Jin H, Kim VN, Cho K-H. The co-regulation mechanism of transcription factors in the human gene regulatory network. Nucleic Acids Res. 2012, 40(18):8849–8861.

82. Gotea V, Visel A, Westlund JM, Nobrega MA, Pennacchio LA, Ovcharenko I. Homotypic clusters of transcription factor binding sites are a key component of human promoters and enhancers. Genome Res. 2010, 20(5):565–577.

83. Terada A, Okada-Hatakeyama M, Tsuda K, Sese J. Statistical significance of combinatorial regulations. Proc. Natl. Acad. Sci. U S A 2013, 110(32):12996–13001.

